# Diverse viruses carrying genes for microbial extremotolerance in the Atacama Desert hyperarid soil

**DOI:** 10.1101/2020.09.21.307520

**Authors:** Yunha Hwang, Janina Rahlff, Dirk Schulze-Makuch, Michael Schloter, Alexander J. Probst

## Abstract

Viruses play an essential role in shaping microbial community structures and serve as reservoirs for genetic diversity in many ecosystems. In hyperarid desert environments, where life itself becomes scarce and loses diversity, the interactions between viruses and host populations have remained elusive. Here, we resolved host-virus interactions in the soil metagenomes of the Atacama Desert hyperarid core, one of the harshest terrestrial environments on Earth. We show evidence of diverse viruses infecting a wide range of hosts found in sites up to 205 km apart. Viral genomes carried putative extremotolerance features (i.e. DNA repair proteins, enzymes against oxidative damage, spore formation proteins) and auxiliary metabolic genes, indicating that viruses could mediate the spread of microbial resilience against environmental stress across the desert. We propose a mutualistic model of host-virus interactions in the hyperarid core where viruses seek protection in microbial cells as lysogens or pseudolysogens, while viral extremotolerance genes aid survival of their hosts. Our results suggest that the host-virus interactions in the Atacama Desert soils are dynamic and complex, shaping uniquely adapted microbiomes in this highly selective and hostile environment.

**Importance:** Deserts are one of the largest and rapidly expanding terrestrial ecosystems characterized by low biodiversity and biomass. The hyperarid core of the Atacama Desert, previously thought to be devoid of life, is one of the harshest environments supporting only scant biomass of highly adapted microbes. While there is growing evidence that viruses play essential roles in shaping the diversity and structure of nearly every ecosystem, very little is known about the role of viruses in desert soils, especially where viral contact with viable hosts is significantly reduced. Our results indicate that diverse viruses are widely dispersed across the desert, potentially spreading key stress resilience and metabolic genes to ensure host survival. The desertification accelerated by climate change expands both the ecosystem cover and the ecological significance of the desert virome. This study sheds light on the complex virus-host interplay that shapes the unique microbiome in desert soils.

## Introduction

Viruses are considered to be the most abundant biological entities on Earth (1), with high genomic diversity (2) and an expanding ecological and biogeochemical importance. Viruses, particularly bacteriophages (thereafter: phages), shape microbial community turnover and composition (3, 4), nutrient cycling (5, 6) as well as microbial evolution (7, 8) in marine (9) and freshwater (10) environments. The progress in soil virome studies is lagging compared to those in marine and gut microbiome systems (11– 13), mainly due to the difficulties in isolating viruses from heterogeneous and complex soil environments (14). However, recent metagenomic approaches revealed diverse soil viruses in high abundance (15), which play significant roles in carbon processing (16–18) and other nutrient turnover (19, 20). Even less explored are viruses in extreme soil environments, where life itself becomes scarce in biomass and low in biodiversity (21–23). Understanding the abundance and diversity of viruses as well as their interactions with extremotolerant microbes in environments can highlight the unique roles viruses may play in driving the adaptation of their hosts, and reveal dispersal and diversification of viruses in sparsely populated and harsh environments.

Hyperarid desert soils are unique terrestrial environments, where low water availability limits proliferation and diversification of life. Biota that permanently inhabits these environments is often limited to a few bacterial and archaeal phyla. Recent studies of warm (i.e. Namib Desert, Sahara Desert) and cold (i.e. Antarctic soil) hyperarid desert viromes have revealed abundant viruses of diverse lineages and sizes, with lysogenic and pseudolysogenic viruses being more prevalent than lytic viruses in hot deserts (21, 24–26). With little water availability and extended periods of drought, hyperarid desert soils present a distinct model for studying viral persistence and dispersal. In these ecosystems, viral mobility is limited compared to aquatic environments, in which both viruses and hosts freely diffuse (14).

The Atacama Desert is one of the harshest environments on Earth with its hyperarid core experiencing extreme desiccation with mean annual precipitation < 2 mm (27). The surface soil of the Atacama hyperarid core generally contains < 1 wt % of water, and experiences high daily ultraviolet (UV) radiation (30 J.m^−2^) (28), extreme diurnal temperature fluctuations (∼ 60°C) (29) and additional osmotic pressure from the accumulation of salts (28, 30). Scarce populations of highly adapted microbial communities consisting of *Actinobacteria, Firmicutes, Chloroflexi* (28, 30, 31) and more recently, *Thaumarchaeota* (29) were found to inhabit soils of the Atacama hyperarid core. However, very little is known about viruses from these desert soil microbiomes. Crits-Christoph *et al*. (32) identified viral sequences and their potential hosts in halite endoliths of the Atacama. Additionally, Uritskiy *et al*. (33) detected transcriptionally active viruses potentially infecting *Halobacteria* also inhabiting halite salt nodules in a salar located in the Atacama Desert. These niche halite host-virus relationships highlight the need to characterize the impact of viruses in broad desert soils that represent one of the largest and rapidly expanding terrestrial ecosystems on the planet (∼35% of the Earth’s land surface (34)). In our previous study (28), we also detected viruses in the Atacama Desert soils of the hyperarid core using read-based analyses of metagenomes, however, no information exists regarding host-virus interactions, dispersal and potential function of these viruses.

To understand the diversity and ecological impact of viruses inhabiting the hyperarid soils, we investigated viral genomes assembled from soil metagenomes of the Atacama hyperarid core. We identified host-virus interactions, innate and adaptive host immunity elements, and phylogenetic diversity of viruses across geographically distant sampling locations. We analyzed putative extremotolerance genes and auxiliary metabolic genes (AMGs) found in the predicted viral sequences, providing evidence for a complex trade-off between viral predation and viral delivery of extremotolerance genes to microbes inhabiting harsh hyperarid desert soils.

## Results

### Soil metagenomes of the Atacama hyperarid core feature heterogeneous viromes

We predicted viral scaffolds in eleven assembled metagenomes (4.1 Gbp in total) from three different boulder fields (Lomas Bayas, L; Maria Elena, M; Yungay; Y), which were previously studied for the impact of boulder cover on the soil microbiome, uncovering highly adapted microbes sheltered below the boulders of expansive boulder fields in the Atacama Desert hyperarid core (29). The aforementioned study compared the microbiome in two different soil compartments (Below boulder, B; Control - exposed soil adjacent to Boulder, C), for which we kept the designation consistent in this paper (for the map of sampling locations see **Figure 1**, for analysis workflow see **Figure S1**). In total, 6809 out of 707,509 examined scaffolds were predicted to be viral. After quality and length filtering, we identified 86 viral scaffolds (hereafter referred to as “viral genomes”) with length >10 kb forming 84 viral “populations” (dereplicated at 99% identity). In detail, Virsorter (35) predicted 79 viral genomes, while VIBRANT (36) predicted 37, including 30 overlapping between the two tools. The average length of the predicted viral genomes was 32.7 kbp (± 29.5 kbp), with the longest being 177 kbp and smallest being 10 kbp. The average G+C content of viral genomes was 58.7 % (± 10.3 %) and the average coding density was 91.0 % (± 4.2 %). The viral genomes were of varying quality: 9.30 % “complete”, 10.5 % “high quality”, 17.4 % “medium quality”, 61.6 % “low quality” and 1.16 % “not-determined” according to CheckV (37). From these 84 viral populations, eight were predicted to be lysogenic. An overview of the viral population genomes can be found in **Table S1**.

**Figure 1.**
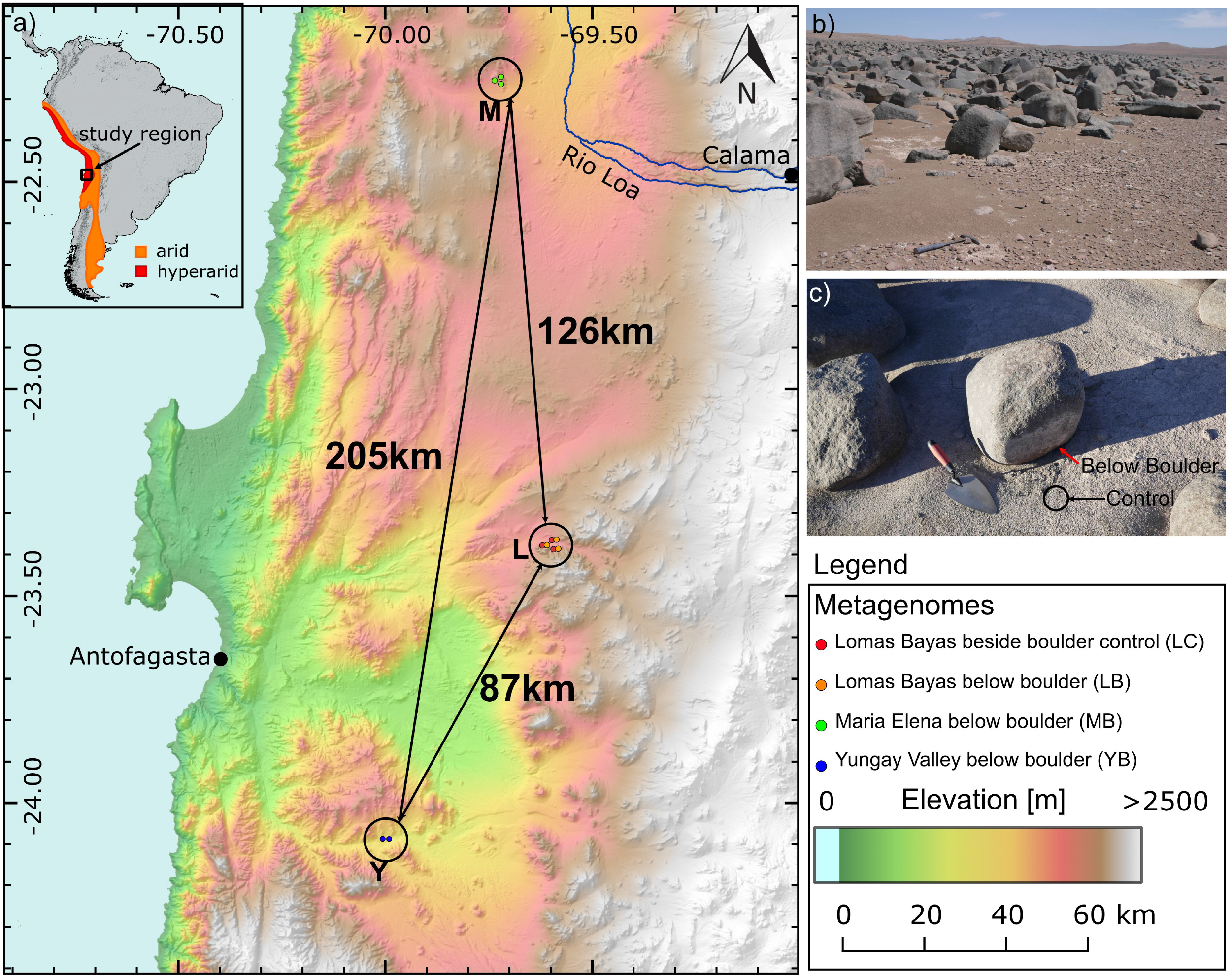
Sampling location information. a) Map of the sampling locations M: Maria Elena, L: Lomas Bayas, Y: Yungay Valley. Distances between sampling locations are shown. Approximate metagenome sample locations are shown in smaller circles. b) Yungay boulder field. c) A sampled boulder in Maria Elena boulder field. Red and black arrows distinguish sample types B and C, respectively.

We observed a large degree of heterogeneity in the viral populations between samples, in terms of both alpha and beta diversity. **Figure 2a** shows the relative abundances of viral populations based on the total number of reads normalized across samples. The mapping-based coverage of viral populations identified in this study varied significantly ranging between 4.5 to 5075, and the total abundance of these viral populations varied up to 65-fold between samples. Notably, we did not detect prevalence of lysogenic viruses in L and M sites, while YB samples exhibited higher relative abundance (∼79%) of lysogenic viruses. Samples collected from L sites (n = 6) had higher alpha diversity and species evenness (**Figure 2b**) than samples from M and Y (n = 5) (Welch’s t-test, p = 0.0022 (alpha-diversity) and p = 0.0056 (species evenness)). Principal Coordinate Analysis (PCoA) (**Figure 2c**) based on Bray-Curtis distances showed clustering by sampling site, while the beta diversities varied between sites, with YB viromes being particularly conserved, and LB and MB viromes exhibiting higher variability compared to LC viromes. Permutational multivariate analysis of variance (PERMANOVA (38)) confirmed statistically significant differentiation of viral communities based on the sampling site (R2 = 0.384, p = 0.011). Biota-environment (BioENV) analysis (39) using geochemical and environmental metadata (**Table S2)** revealed temperature and Na^+^ concentration to be most correlated with the viral community composition (rho = 0.5642, p-value = 0.001). Majority (66.7%) of the viral populations were unique to the sample, and only one lysogenic virus with a representative genome of length 12.6 kbp was detected across all three sites in eight samples, and three additional viral populations were observed between two different sites (**Figure 2d**).

**Figure 2.**
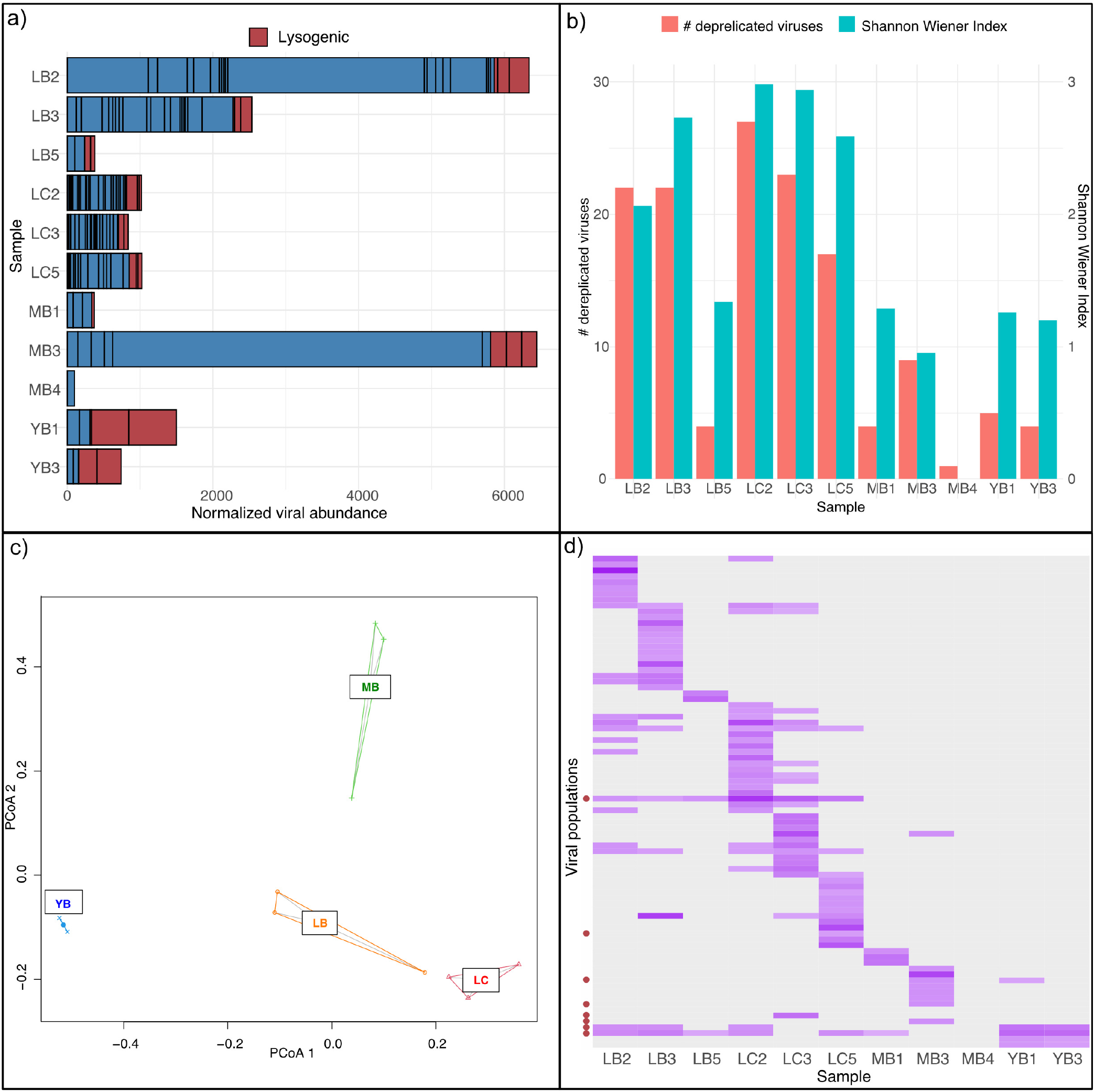
Abundance and diversity analyses of viromes. a) Abundance of viral contigs normalized by sequencing depth of each sample. Each bar denotes a single viral population, and predicted lysogenic viral populations are marked in red. b) Alpha diversity and species evenness of viromes. c) PCoA plot of viromes using Bray-Curtis distance matrix. d) Relative abundance profile of each viral population where darker purple denotes a viral population with highest relative abundance in a sample (grey means absence). Lysogenic populations are marked with red circles on the y-axis. Graphs were visualized in R v4.0.2(81) using ggplot2 (83)

By comparing the distributions of viral and microbial genome abundances in each sample (**Figure S2**), we found that only LC samples featured statistically significantly lower abundances (p-value < 0.05, Welch’s t-test) in the viral genomes compared to microbial genomes, while no statistically significant differences could be observed in other samples. However, the ranges of viral genome abundances were greater than those of microbes in all samples, with LB2 and MB3 samples featuring viruses with abundances up to two orders of magnitude greater than those of microbes. We conducted symmetric co-correspondence analysis (sCoCA) to test whether the viral and microbial community compositions covary. The best sCoCA model using the first three axes determined the common variance between the viral and microbial communities to explain 37.2% and 56.8% of the total variances of viral and microbial communities, respectively (p = 0.006). The first three axes computed by sCoCA accounted for the 62.3% (CoCA1 = 26.5%, CoCA2 = 20.4%, CoCA3 = 15.3%) of the common variance. The first three axes of the ordinations in viral and microbial communities were highly correlated with each other (Pearson product-moment correlation coefficient > 0.99). The relative abundances of microbial taxa across samples used to conduct sCoCA are visualized in **Figure S3a** and the ordination biplots (**Figure S3b-c**) illustrate highly similar positioning of the samples along the first two axes identified by sCoCA between microbial and viral communities. Higher species richness and species evenness (Shannon indices) could be observed in LB and LC samples for both microbial and viral communities, however, little pattern in the microbial population could be observed at the phylum level, suggesting the covariance patterns to be rooted in the abundance profiles of individual populations. Interestingly, neither viral nor microbial communities were predictive of each other when conducting CoCA in the predictive mode (p > 0.05).

### Atacama viruses are phylogenetically novel and diverse

We clustered 84 dereplicated viral population genomes using intergenomic similarities (40). 82 clusters were formed at the genus level (intergenomic similarity threshold at 70%), indicating all except two viral population genomes recovered in this study to be of different genera. The OPTSIL clustering of viruses (41) yielded 84, 64 and 14 clusters at the species, genus and family level (**Figure S4**). vConTACT2 (42) was used to cluster the Atacama viral genomes with 2616 known prokaryotic viruses (**Figure 3a**). Twenty-two Atacama viral genomes were related to phages infecting *Gordonia, Mycobacterium, Streptomyces* and *Arthrobacter* at a taxonomic level higher than genus. Only four viral contigs were predicted to be in the same genus as a *Gordonia* phage, a *Streptomyces* phage, a *Lactococcus* phage and an *Arthrobacter* phage from the reference database, all belonging to the order *Caudovirales*, family *Siphoviridae* (tailed dsDNA phage). vConTACT2 also predicted 20 genus-level clusters consisting of two to five Atacama viruses (Atacama Viral Clusters [AVC]). Of the remaining viral populations that could not be clustered, 27 were classified as “singleton” viruses that share no or very few genes with the database and each other, and 19 “outlier” viruses that could be associated with existing sequences but could not be clustered due to low confidence level. Atacama viral genomes tended to cluster with each other in the gene sharing network rather than with the viral sequences in the database. Notably, the majority of the Atacama viruses that are related to reference *Streptomyces* phages and *Mycobacterium* phage were recovered from the LC samples. Across the three different tools, we estimated between 64 and 80 genera-level clusters amongst the 84 viral populations.

**Figure 3.**
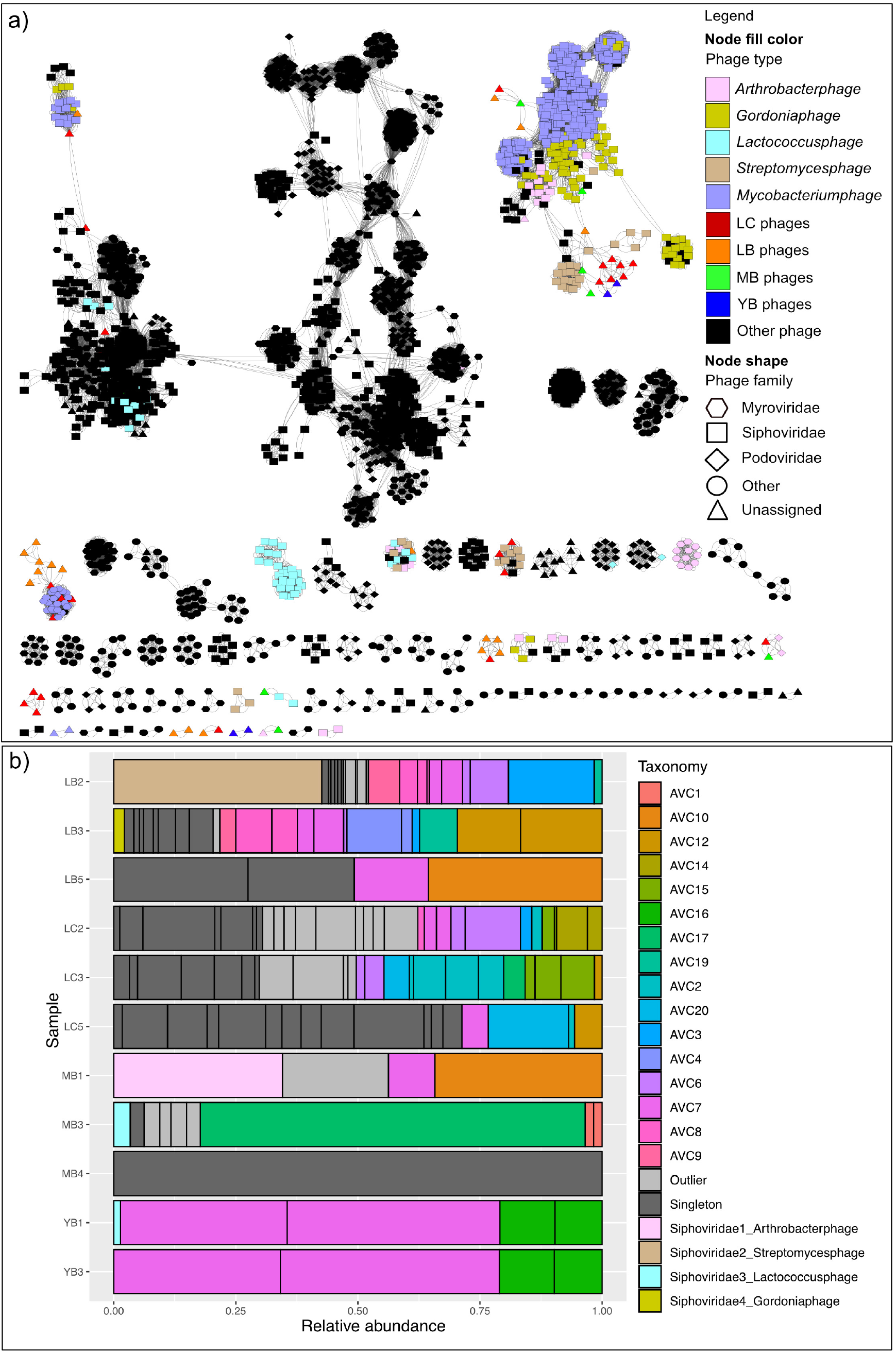
Clustering of Atacama viral genomes with reference viral genomes based on shared genes. a) vConTACT2 output network with significance greater than 1. Network was visualized using Cytoscape v3.8.0 (90). Queried viruses are color coded based on the sampling site they were recovered from and reference viruses are color-coded based on the host they were annotated to infect. Node shape denotes the phage family of reference viruses. b) Relative abundance of identified taxonomic groups per sample. AVC denotes genus-level Atacama Viral Clusters identified using vConTACT2. Each bar represents a distinct viral population.

The relative abundance profiles of different taxonomic groups in each sample (**Figure 3b**) illustrated a high level of heterogeneity between the three sites as well as amongst the samples collected from sites L and M. In contrast, the two Y samples were almost identical in the taxonomic composition of their viromes. For all the samples, singletons and outlier clusters of viral genomes that are biologically novel (due to very little protein homology to the existing database) consisted the majority.

### Sequence-informed putative host-virus interactions indicate dispersal of hosts and/or viruses

Previously, we recovered 73 medium to high quality (>75% completeness; <15% contamination) MAGs across eleven metagenomes from three sampling locations (29). The MAGs were classified as 34 *Actinobacteria*, 30 *Chloroflexi*, eight *Thaumarchaeota* and one *Firmicutes*. We resolved 74 unique interactions between 30 MAGs and 15 viruses using the following four sequence-based methods: 1) protospacer-to-spacer match for MAGs containing CRISPR arrays, 2) oligonucleotide frequency similarity (VirHostMatcher (43)), 3) tRNA matching and 4) nucleotide sequence homology. **Figure 4a** illustrates the putative host-virus interactions specifying the method, with which the interactions were resolved.

**Figure 4.**
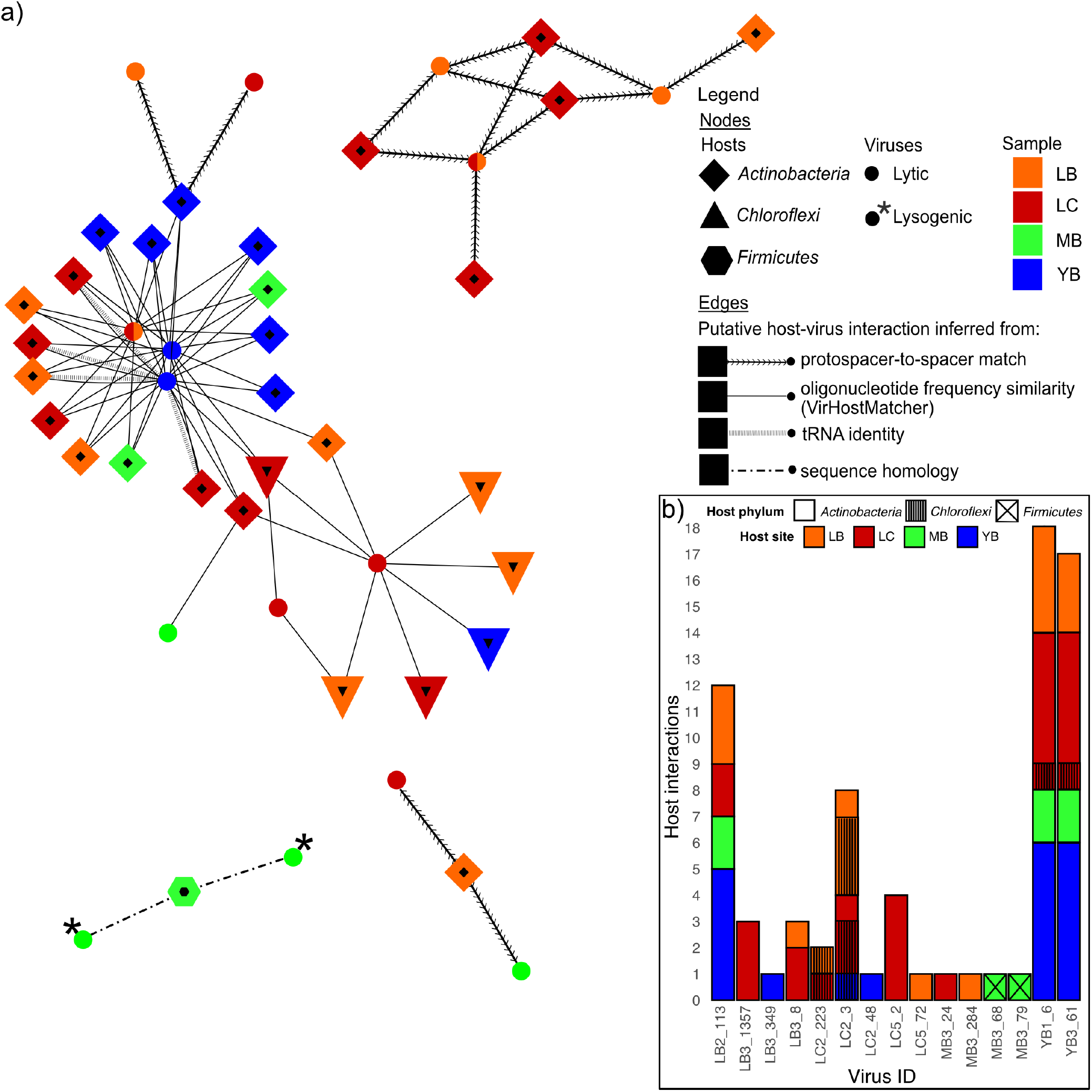
Host-virus interactions. a) Putative host-virus interaction network visualizes host taxonomy, host location, virus location (if a virus was detected in multiple samples, multiple colors are used) and virus lifestyle. Edges denote the method used to predict the host-virus interaction. Visualization was done using Cytoscape v3.8.0 (90). b) Number of host interactions per virus, where the bar color denotes the site from which the matched MAGs were assembled and bar texture denotes host taxonomy.

We identified 14 unique interactions between six actinobacterial MAGs with CRISPRs and seven viruses. Out of 73 MAGs surveyed, nine actinobacterial and two Chloroflexi-derived MAGs contained CRISPR arrays from which direct repeat (DR) sequences were extracted. All MAGs carried unique sets of DR sequences, although three identical DRs appeared across six MAGs (see **Table S2** for detail). In total, 18 identified DR sequences were used to recover 3438 unique spacers directly from the reads in their respective metagenomes. Five actinobacterial MAGs classified as *Rubrobacters* recovered from site L clustered based on their shared infection histories with three phylogenetically distinct viruses also detected from site L, assuming that the CRISPR arrays were not horizontally transferred (44). Interestingly, two additional actinobacterial MAGs (one *Rubrobacter*, one *Acidimirobiia*) had acquired spacers that results in resistances against viruses detected from sites between 87 and 205 km away (**Figure 2**), which indicates viral and/or host dispersal across the desert. In sum, spacer to protospacer-based identification of host-virus interactions revealed potentially widely dispersed viruses preying on *Actinobacteria*, particularly *Rubrobacters*.

Although the spacer to protospacer matches between CRISPR containing MAGs and viral genomes provide high confidence evidence of historical infections between a host population and viruses, many bacteria and archaea do not have CRISPR-*Cas* defense systems (45) or the respective CRISPR arrays do not get assembled or binned into MAGs. In the studied metagenomes, only 17% of the medium to high quality MAGs contained CRISPR arrays. To predict possible host-virus interactions for hosts that lack CRISPR systems, VirHostMatcher (46) identified 54 putative interactions between 23 MAGs (17 *Actinobacteria*, six *Chloroflexi*) and six viral genomes based on shared k-mer frequency patterns. Most linkages were established between 16 actinobacterial MAGs belonging to class *Acidomicrobiia* and three viruses belonging to two genus level clusters. Interestingly, VirHostMatcher matched some viruses to hosts that are taxonomically distant, some differing at the order level and a few even in different phyla. For instance, three viruses matched to both *Chloroflexi* and *Actinobacteria* (**Figure 4b**) suggesting the possibility that these viruses have broad host ranges. No overlaps between interactions based on spacer-to-protospacer matches and VirHostMatcher host-virus linkage were identified.

Additionally, tRNA matching and sequence homology search were conducted. We detected between one and 35 tRNAs across 20 viral population genomes. Only complete identity between viral and host tRNAs were used as an indication of potential host-virus interaction. We identified four interactions between four actinobacterial MAGs and one virus detected from the Y site. These interactions overlapped with interactions inferred from oligonucleotide frequency similarity. Finally, nucleotide sequence homology was used to identify putative host-virus interactions between two lysogenic viruses and a firmicutal MAG. **Figure 4b** summarizes the putative host-virus linkages for each viral genome, visualizing a high degree of variance in the number of identified interactions per virus, as well as the evidence of putative cross-site host-virus interactions and the potential for broad host ranges for some viruses.

### Atacama viruses encode genes against environmental stress

Across 84 viral populations, we identified and annotated 4,288 proteins using DRAM-v (47). Thirtynine percent of these proteins could be associated with the sequences in the queried database with high confidence, of which approximately half were “uncharacterized” and “hypothetical” proteins. We found 93 genes likely involved in extremotolerance of microorganisms (for loci information see **Table S4**) across 40 viral populations. For instance, numerous DNA repair proteins (double-strand break repair protein, RecA, RecT, resolvases, UvdE, ribonucleotide reductases, thymidylate synthase, oxidoreductase, dUTPase) were identified, which could provide resistance against oxidative stress in the host. Additionally, we found a sporulation protein (Spherulation-specific family 4 protein, SSF) in three viral genomes assembled from the LC2 sample. Other putative AMGs included PhoD-like phosphatase, esterase, glucanases, glycosyl hydrolases and endo-beta-N-acetylglucosaminidase. Some viruses also encoded membrane transport proteins for cation transporters and potassium channels, while others encoded transcriptional factors including WhiB (48), which is also involved in bacterial sporulation initiation (49). Interestingly, we identified LuxR among the AMGs, the response regulator involved in quorum sensing of bacteria, which binds homoserine lactones and activates genes in the respective operon (50). Extremotolerance genes and other AMGs were present across all sampling sites (**Figure 5a**), with the LB site exhibiting the highest abundance of viruses carrying these genes, while the LC site harboring the highest diversity of these genes. **Figure 5b** visualizes genomic regions of example scaffolds carrying AMGs, displaying relatively uniform coverage across all gene loci and a close vicinity of viral hallmark genes and AMGs providing direct evidence that these are bona fide AMGs.

**Figure 5.**
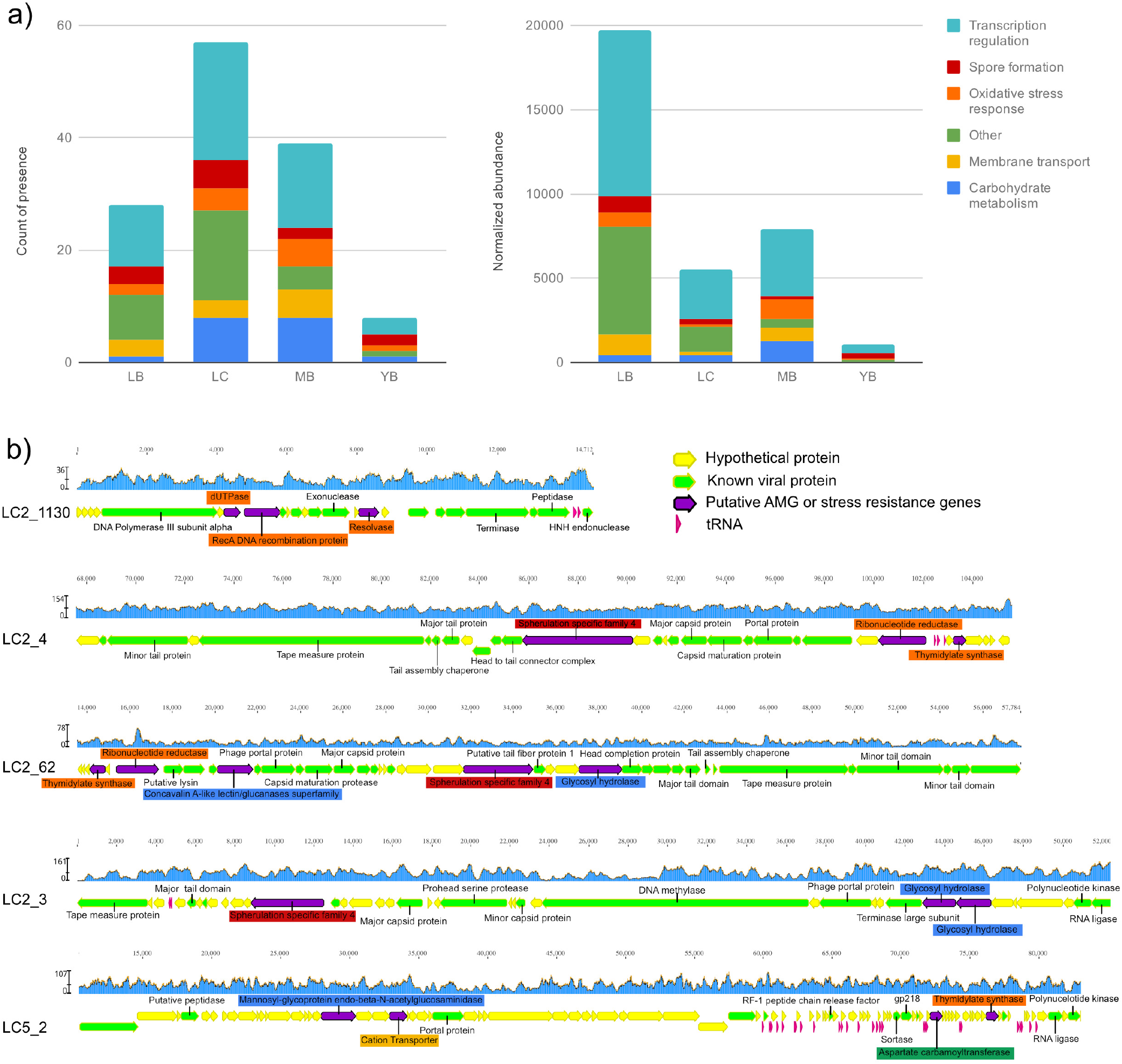
Extremotolerance and auxiliary metabolic genes. a) Sums of count of presences (left) and normalized abundance (right) of putative extremotolerance genes and AMGs across sampling sites. Abundances were calculated using the coverage of scaffolds where the gene was found in and normalized across samples by sequencing depth. b) Genomic neighborhoods of putative extremotolerance genes and AMGs. Genes of interest are colored purple and labels are color-coded by the functional categories according to the color legend used in a). Identified viral hallmark genes are labeled and colored green, while genes homologs to a viral protein in the DRAM-v (47) database without a known function are colored green without a label. Hypothetical genes and tRNAs are denoted in yellow and pink respectively. Coverages calculated based on mapped reads are shown in the blue graph above each genomic region visualized with Geneious v11.1.5.

We further investigated the evolutionary relationship of two putative viral AMGs relative to host AMGs, SSF genes and *whiB*. These two genes are generally known to be associated with bacterial spore formation and therefore could provide key extremotolerance for microbes in harsh desert environments. We compared the three genomic regions containing the SSF genes (**Figure S5a**), and found that the genes upstream and downstream the SSF genes in the viral scaffolds LC2_4 and LC2_62 showed synteny despite the lack of sequence homology between the SSF proteins themselves. On the other hand, SSF proteins found in LC2_62 and LC2_3 contained homologous regions, despite the lack of synteny and the disparity in the GC content throughout the genome. We examined the evolutionary history of these SSF proteins by placing them on a phylogenetic tree with similar bacterial proteins deposited in the NCBI nr database (**Figure S5b**) and ten SSF protein homologs found across nine MAGs from this environment. Interestingly, only actinobacterial MAGs binned from the L site metagenomes contained this gene and formed two distinct clusters with each other. Compared to publicly available SSF proteins, the viral SSF proteins identified herein were not closely related to their homologs found in reconstructed MAGs. The closest homologs to the viral SSF proteins were found in genomes of *Actinobacteria* (e.g., *Streptomyces*), which contains many spore-forming bacteria. Notably, the viral genome of LC2_3 was predicted to have interactions with *Actinobacteria* and *Chloroflexi* in our samples (based on oligonucleotide frequency similarity; **Figure 4**), and two actinobacterial MAGs also encoded distantly related SSF proteins (predicted virus-MAG interactions are shown with red lines in **Figure S5b**). Based on the evidence of unique SSF proteins found in three distinct viral genomes and the phylogenetic divergence from host proteins, the likelihood that they are randomly packaged host genes is low. Although the function of the viral SSF proteins requires experimental confirmation, we suspect this relatively large gene (mean protein length 1265 amino acidss) to be beneficial for the viral populations, as it was identified in multiple viruses despite the presumed high cost of maintenance of AMGs in viral genomes (51, 52).

WhiB-like transcription factor has previously been shown to control the spherulation septation in *Actinobacteria* (49). In addition, WhiB-like proteins previously identified in several *Mycobacterium* phages and *Streptomyces* phages (53) have been shown to regulate host cell wall component alteration in *Mycobacteria* (54). Twelve WhiB-like transcription factor genes were identified across eleven viral populations as well as in 49 actinobacterial MAGs across all three sites. We phylogenetically placed the twelve viral WhiB-like proteins with bacterial WhiB proteins found in medium-to-high quality MAGs (**Figure S6a**). Viral WhiB-like proteins were generally more related to each other than to the bacterial proteins. Additionally, WhiB-like proteins found in interacting virus-microbe pairs (**Figure S6a**) were not closely related compared to other homologs. Notably, viral WhiB-like proteins from the same site tend to cluster together phylogenetically, with the exception of the LC samples. When comparing these proteins with related sequences in the NCBI nr database, we found four clusters with phage proteins, while the rest (eight proteins) clustered with homologs found in bacteria (**Figure S6b**).

## Discussion

The hyperarid core of the Atacama Desert harbors an abundant and diverse soil virome that remained mostly understudied due to the scarcity of microbial biomass available. Recent improvements in soil DNA isolation methods and deeper sequencing of metagenomes (28, 29) not only allowed the discovery of microbes actively replicating *in situ*, but also shed light upon the viral fraction of the hyperarid soil ecosystem that coexist with their microbial hosts. Our investigation of the Atacama viromes reveals taxonomically diverse viruses and complex interactions between viruses and their hosts across the desert. Notably, the viruses contained key extremotolerance genes, and we propose a mutualistic model of host-virus interaction, where viruses seek protection in microbes as lysogens and pseudolysogens and in return, aid host extremotolerance and survival.

### Diversity of Atacama viruses contradicts the scarcity of microbial hosts

The Atacama Desert soil virome analyzed in this study consists of 84 viral populations belonging to at least 60 novel genera. The diversity of the viruses in the Atacama Desert soils are astonishing considering that the microorganisms that inhabit these soils are low in both biodiversity and biomass. Typically, groups of viruses infecting the same host often exchange genetic materials, and form genotypic clusters (55). Therefore, in a low diversity ecosystem, where many viruses infect the same hosts, one may expect a stronger genotypic clustering of viruses. Additionally, large diversity of viral predators coupled with low diversity of prokaryotic prey seems to go against the ‘competitive exclusion principle’ (56). High abundance and diversity of viruses in an environment with reduced encounters with viable microbial hosts suggest that some of the Atacama soil viruses may be dormant virions waiting for the appropriate host population to thrive, while others remain protected by residing in the host cells as lysogens (integrated into host genomes or plasmids) or pseudolysogens (as virus particles in the host cytoplasm) (24, 57). In particular, viruses have been shown to seek protection in their host cells and this mode of viral survival has been observed in hot hyperarid desert soils, where lysogenic and pseudolysogenic (hereafter referred to as “(pseudo)lysogenic”) phages were found to be more prevalent (24–26). In our study, we predicted eight viral “populations” to be lysogenic based on their representative genomes being proviral. However, Computational prediction tools tend to underestimate lysogenic viruses (35) and cannot distinguish pseudolysogens from lytic viruses. Therefore, isolation, cultivation and visualization approaches of the viruses and their hosts would shed light on the lifestyle of the viruses in the Atacama Desert hyperarid soils once sufficient biomass can be harvested from the ecosystem.

### Viral dispersal is likely host-mediated across the Atacama Desert

Our study demonstrated high co-correspondence between the viral and microbial communities, possibly due to the sample and site-specific environmental stressors controlling both microbial and virus populations. Despite the high heterogeneity and specificity between samples, we identified four viral populations (two of which were lysogenic) detected in two or more sites (L, M, Y). None of such genomes could be linked to a host, while 35 out of 74 predicted host-virus interactions were between microbes and viruses detected at different sites. In particular, four host-virus interactions based on protospacer-to-spacer matches provide evidence for past dispersal events of either microbial and/or viral populations across distances up to 205 km. We hypothesize one specific mechanism of dispersal in the Atacama Desert to be the frequent sandstorms and powerful winds (58, 59) transporting infected microbes and virions in organic aerosols (60, 61). Notably, the most frequently detected viral population across samples (in eight out of eleven samples) was predicted to be lysogenic, supporting the scenario of viral dispersal through the transport of infected hosts. Additionally, we observed statistically significant lower abundance of viral entities relative to microbes (**Figure 5S**) in C samples suggesting that viruses are vulnerable to irradiation in desert environments, perhaps more so than microbes. This result may also indicate that viral entities sheltered below boulders are undergoing lytic cycles and therefore exhibiting higher abundance relative to microbes, while those beside boulders are primarily (pseudo)lysogenic. Future work is required to reveal the predominant lifestyle and viability of viruses in the Atacama hyperarid core.

### A mutualistic model of host virus interactions in the Atacama Desert hyperarid core

A closer look at the genes carried by the Atacama viruses suggested an intriguing interaction between viruses and hosts, where a fine balance between viral predation and host extremotolerance sustains the continuum of the ecosystem. We posit that the viruses may serve as vectors delivering extremotolerance genes to their microbial hosts, increasing the chance of microbial survival under harsh conditions of the Atacama Desert hyperarid core. In particular, we propose a model specific to extreme deserts, where (pseudo)lysogenic viruses encoding extremotolerance genes could support microbial survival in exchange for taking up shelter inside the bacterial cytoplasm or genome. This mutualistic model parallels well with viral AMGs found in temperate environments (i.e. photosystem I and II genes in marine cyanophages (62–64), CAZYmes in mangrove soil viruses (65)), where AMGs are selected to maximize viral production by enhancing host metabolism during an infection. In the hyperarid core of the Atacama, viral extremotolerance genes likely increase the chance of viral production by aiding host survival, even if they result in temporary dormancy of the host through sporulation (in the case of SSF protein and WhiB). A similar model of virus-host interactions (66) has been described in biofilms, where lysogenic viruses support formation, stabilization, and dispersal of biofilms, and biofilms in return provide protection for viruses against environmental stress (67–69). For instance, in hot desert soils, where microbes are known to form biofilms to protect themselves from desiccation, UV radiation, and poor nutrient availability (70), Zablocki *et al*. (21) hypothesized a positive selection for temperate viruses in biofilms. In even more extreme desert environments such as the hyperarid core of the Atacama, where we did not detect any formation of biofilms, viruses may instead seek protection in the cytoplasm of the microbes as (pseudo)lysogens as previously identified in other hot deserts (24–26). In this case, viral genes encoding extremotolerance may be selected for two reasons: 1) to ensure the survival of both the microbe and the (pseudo)lysogenic virus in the short-term and 2) to spread extremotolerance genes amongst microbes via transduction or lysogeny, resulting in the long-term increased fitness of the hosts against environmental stress. A visual schematic of the proposed host-virus interactions in the Atacama hyperarid core and comparison with proposed host-virus interactions in other environments are found in **Figure 6**. This mutualistic model does not exclude the antagonistic interactions between viruses and their hosts, as evidenced by the diverse innate and adaptive antiphage systems encoded in host MAGs (for analysis on innate and adaptive immune systems please see **Supplementary Results and Discussion**). We posit that viruses undergo lytic cycles when more favorable environmental conditions are met (i.e. rain events) or even in sheltered environments, as evidenced by higher relative viral abundances under boulders compared to beside boulders. Phylogenetic analyses of the viral homologs to SSF proteins and the WhiB-like transcription factors suggest that these genes are indeed distinct from bacterial homologs, including those found in putative hosts. We also observed phylogenetic clustering of viral homologs to WhiB-like transcription factors by sampling site, suggesting the possibility of individual adaptation to the host communities and/or the environmental conditions of each sampling site. Model host-virus systems will be required to confirm our proposed model (**Figure 6**) and identify to which extent extremotolerance genes provide an increase in fitness for microbes and viruses in hyperarid environments.

**Figure 6.**
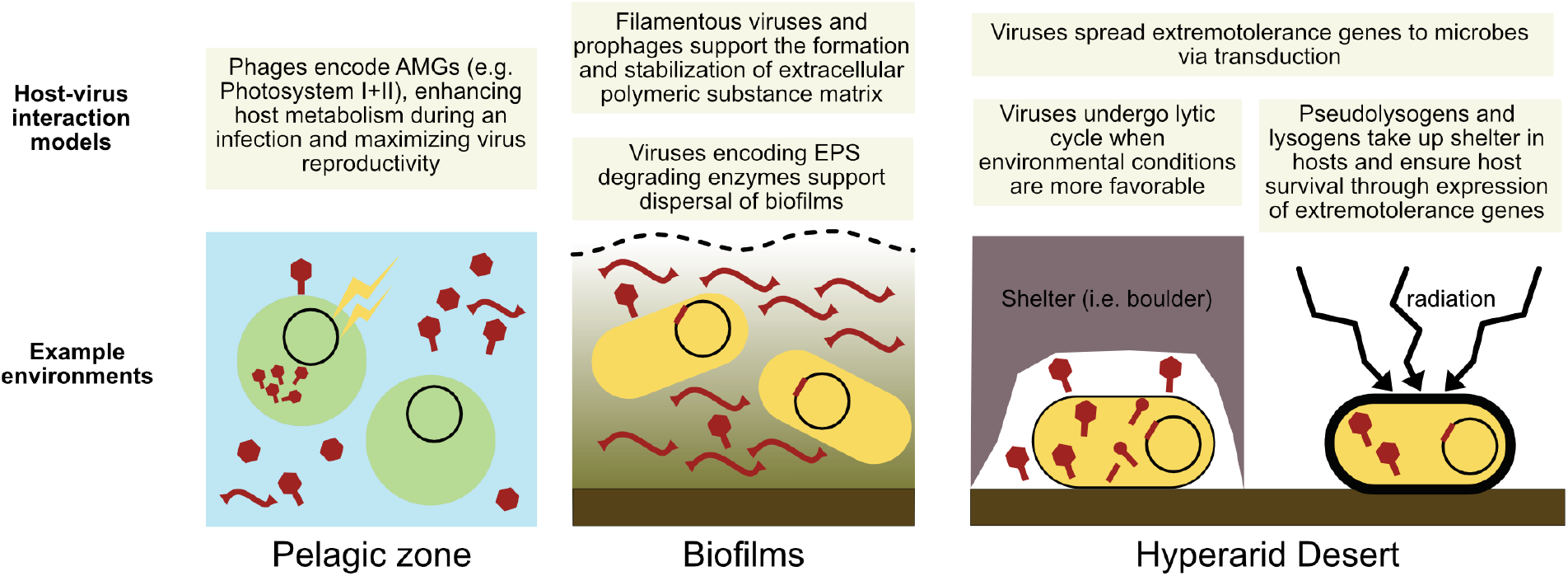
A visual schematic of the proposed host-virus interaction models in the hyperarid core of the Atacama Desert and comparison with models in other environments. Viruses (virions, pseudolysogens and prophages) are shown in dark red, and microbial cells in yellow and green.

### Conclusion

The hyperarid core of the Atacama Desert is a much more biologically complex ecosystem than previously thought. We investigated hyperarid soil metagenomes to uncover a diverse virome interacting with a wide range of microbial hosts. Viruses in the Atacama Desert not only endure long periods of desiccation and extreme oxidative stress themselves, but may also deliver extremotolerance genes to their hosts and aid their survival. This study expands the ecological significance of viruses in terrestrial systems, particularly in deserts. Life seems to persist even in the most hostile environments on Earth, and so do viruses. The Atacama Desert virome and its complex interplay with the extremophilic host populations highlight the role viruses play in microbial evolution and dynamics and propose a new dimension to host-virus interactions in extreme environments.

## Materials and Methods

### Sampling location, procedure and metagenomic library preparation

Briefly, sampling was conducted in March 2019. Three sampling sites, Yungay (Y), Maria Elena (M), and Lomas Bayas (L), were chosen from the hyperarid core of the Atacama Desert (map of the sampling locations shown in **Figure 1**). Samples were collected from below boulders (B) and in the exposed surface soil (control; C) beside boulders. Three B samples (LB2, LB3, LB5) were collected at the Lomas Bayas boulder field, three B (MB1, MB3, MB4) samples from the Maria Elena boulder field and two B (YB1, YB3) samples from the Yungay boulder field. Each B sample was collected from soil below one unique boulder, where the number in the sample name corresponds to the specific boulder. Three C (LC2, LC3, LC5) samples were taken from the Lomas Bayas boulder field, from the exposed topsoil beside corresponding sampled boulders. Eleven metagenomic libraries of DNA extracted from eight B samples and three C samples were sequenced on Illumina HiSeq 2500 (illumina, CA, USA). Detailed sampling procedure, site coordinates, DNA extraction and Illumina library preparation and sequencing can be found in our previous study(29).

### Metagenomic analysis, host genome binning and taxonomic classification

Assembly of metagenomic reads, contig binning, and bin analyses can be found in our previous study (29). Only medium-to-high quality bins (>75% completeness and <15% contamination calculated using CheckM v1.0.13 (71)) were considered as host genomes. Host taxonomy was predicted using GTDB-Tk classify_wf (72). MAGs underwent gene prediction using Prodigal (73) in meta mode and annotated using Diamond v0.9.9 (74) against the UniRef100 database (75). The annotations were subsequently screened for identification of host innate defense marker genes identified by Beziudt *et al* (76).

### Prediction and analysis of viral scaffolds

A schematic illustration of analyses conducted can be found in **Figure S1**. VirSorter v1 (35) with default settings and –diamond flag, as well as VIBRANT v1.2.1 in default settings (36) were used for viral signal prediction across all assembled metagenomes and scaffolds length >=1000 bp. Virsorter predicted viral scaffolds in categories 1 and 2 were combined with VIBRANT predicted viral scaffolds with qualities medium, high and complete. A viral contig was considered to be lysogenic if predicted so by at least one of the following tools: VIBRANT, VirSorter and CheckV v0.6.0. Similarly, a virus was considered complete if at least one of the three previously mentioned tools predicted it to be circular or complete. Viral contigs were dereplicated using CD-HIT v4.6 (77) at 99% identity to identify viral populations, which were used for all subsequent analyses. CheckV v0.6.0 (37) was used for completeness and quality estimation. Abundances of viral genomes were estimated using coverage calculated across all samples using a method described by Roux *et al*. (78). In short, the number of reads mapped using Bowtie2 (79) at ≥90% identity using options --ignore-quals –mp = 1,1 –np = 1 –rdg = 0,1 –rfg = 0,1 --score-min = L,0,-0.1 as suggested by Nilsson *et al*. (80). Coverages were calculated only for scaffolds with mapped reads across ≥75% of scaffold length with ≥1 × coverage, for which average per-base coverage was calculated. Abundances of MAGs were estimated using the coverage of ribosomal protein S3 (rpS3) containing scaffold as described in our previous study (29). Calculated coverages were then subsequently normalized across samples by the total number of reads per sequenced library, to control for the sequencing depth of each sample. Statistically significant differences between viral and microbial abundance per sample were determined using Welch’s t-test with p-value threshold of < 0.05. Viral genome annotation was performed using DRAM-v (47) using the UniRef90 database (75) and all other default databases in DRAM except the KEGG annotations.

### Statistical characterization of viromes

Following community statistical analyses based on the normalized coverages of viral populations across samples were conducted and visualized in R version 4.0.2 (2020-06-22) (81) using the Vegan (82) and ggplot2 (83) packages respectively: Shannon-Wiener indices, Bray-Curtis distance matrices (84) calculation and subsequent Principal Coordinate Analysis (PCoA), PERMANOVA (using ‘adonis’ package, (38)), Co-correspondence analysis (CoCA) (using ‘cocorresp’ package, (85)) and BioENV analysis (39). Previously reported (29) environmental and geochemical variables (**Table S2**) were used as input for BioENV. CoCA was performed on microbial and viral relative abundance profiles using the cocorresp package in symmetric mode. Significance and degree of covariance were computed and the ordination biplots were visualized using a method described previously by Alric *et al*. (86). All permutation-based tests were conducted with 999 iterations. Microbial relative abundance profiles were calculated using read-mapping based abundances of *rpS3* genes from our previous study (29).

### Intergenomic distance clustering and phylogenetic analysis of putative viruses

Intergenomic distances of viruses were calculated to identify genus and species level clusters using VIRIDIC with default settings (87). Phylogenetic trees were constructed using nucleic acid sequence-based VICTOR (41). vConTACT2 v.0.9.19 (42, 88) was used to cluster and classify selected viral scaffolds against the ProkaryoticViralRefseq v94 database (89), resulting clusters were subsequently visualized using Cytoscape v3.8.0 (90). Per sample relative abundances of viral clusters were calculated by summing up the calculated coverages for each viral population genome in the cluster and then dividing it by the total coverage of all viral population genomes in the respective sample.

### CRISPR-Cas analysis and spacer extraction of medium-to-high quality host genomes

For each high quality host genome, direct repeats and *Cas* genes associated with CRISPR systems were extracted by combining the tools PILER-CR v1.06 (91) in default settings and CRISPRCasFinder (92) with results filtered for evidence level 4 for the latter. Filtered direct repeats were subsequently used for spacer extraction using MetaCRAST (93) with flags -d 3 -l 60 -c 0.9 -a 0.9 -r from the raw reads of the respective metagenome the MAG was binned from. Spacers were dereplicated using CD-HIT v4.6 (77) at 100% identity to identify the number of unique spacers across all metagenomes and within each sample.

### Host-virus matching

1) Protospacer-to-spacer matching: Extracted CRISPR spacers from all metagenomes were BLAST-ed (94) with blastn --short algorithm against the predicted viral sequences across all metagenomes and filtered with an 80% similarity threshold, (similarity=alignmentLength*Identity)/QueryLength). 2) Oligonucleotide frequency based matching: VirHostMatcher (43) was used to determine putative interactions between medium-to-high quality MAGs from all metagenomes and predicted viral genomes based on shared oligonucleotide frequency pattern (k = 6). A d_2_* dissimilarity threshold of < 0.2 was used to filter all potential host-virus interactions based on the benchmarking performed by Ahlgren *et al*., (2017) where the lowest dissimilarity score threshold of 0.2 yielded above 90% accuracy in host prediction at the class level and approximately 60% accuracy at the order level. 3) tRNA identity based matching: tRNAs in viral genomes and MAGs were predicted using DRAM-v (47) and tRNAscan-SE (95) respectively. Viral tRNAs were BLAST-ed (94) against microbial tRNAs and only complete (100% identity) matches were considered to infer host-virus interactions. 4) nucleotide sequence homology based matching: viral genomes were BLAST-ed against medium-to-high quality MAGs with cut off ≥75% coverage over the length of the viral contig, ≥ 70% minimum nucleotide identity, ≥50 -bit score, and ≤0.001 e-value. Exact matches resulting from a viral genome being binned in a MAG were excluded as potential binning errors, except when the viral genome was identified by Virsorter to be a defined prophage region inside a longer scaffold. Identified interactions were combined and the host-virus interaction network was visualized using Cytoscape v3.8.0 (90).

### Genomic neighborhood visualization and phylogenetic analyses of viral genes

Genomic neighborhoods were compared and visualized using Geneious v11.1.5. Software (https://www.geneious.com) and Easyfig v2.2.2 (96) with BLASTn e-value threshold of 1E-5. Three viral spherulation-specific proteins and twelve viral WhiB-like protein amino acid sequences were queried against the NCBI nr database using BLASTp. Twenty highest identity matches per spherulation-specific sequence and five highest identity matches per WhiB-like sequence were selected for subsequent phylogenetic analyses. Spherulation-specific proteins were searched for in the medium-to high quality MAGs using hmmsearch (HMMER v3.2, www.hmmer.org) with spherulin4.hmm with an e-value threshold of 1E-10. Bacterial WhiB-like protein sequences were selected from the medium-to-high quality MAGs based on their annotation against the UniRef90 database (75). Duplicate sequences were removed prior to alignment using MUSCLE v3.8.31 (97). Spherulation-protein alignments were trimmed using BMGE v1.12 (98) due to the presence of larger gaps. Trees were constructed using iqtree v1.5.5 (99) with flags -m MFP - alrt 1000 -bb 1000 and visualized using iToL (100). Branches with bootstrap (101) values at least 95 and SH-aLRT(102) test values at least 80 were marked as strongly supported. Phage promoters were predicted using PromoterHunter from phiSITE (103). Rho-dependent and Rho-independent terminators were predicted using RhoTermPredict (104) and ARNold (105) respectively.

## Data availability

MAGs and viral genomes used in the analyses are deposited to NCBI under Bioproject PRJNA665391. NCBI accession information for the viral genomes and MAGs are found in **Table S1 and S5**, respectively.

## Acknowledgements

This work was funded by ERC Advanced Grant HOME (# 339231) to DSM. AJP was supported by the Ministerium für Kultur und Wissenschaft des Landes Nordrhein-Westfalen (“Nachwuchsgruppe Dr. Alexander Probst”). We acknowledge support by the German Aerospace Center (DLR) under contract DISPERS (50WB1922). We thank Till Bornemann for providing the script for viral contig coverage calculations.

## Author contributions

YH, AJP and DSM conceived the project; YH conducted sampling; MS generated the raw sequence data; YH assembled, curated and analyzed sequence data with contribution from JR and AJP; AJP provided computational resources; YH wrote the manuscript with contribution from JR; all authors discussed and revised the manuscript.

